# Roosting in exposed microsites by a nocturnal bird, the rufous-cheeked nightjar: implications for water balance under current and future climate conditions

**DOI:** 10.1101/206300

**Authors:** Ryan S. O’Connor, R. Mark Brigham, Andrew E. McKechnie

**Affiliations:** DST-NRF Centre of Excellence at the Percy FitzPatrick Institute, Department of Zoology and Entomology, University of Pretoria, Private Bag X20, Hatfield 0028, South Africa.; Department of Biology, University of Regina, Regina, Saskatchewan S4S 0A2, Canada..; South African Research Chair in Conservation Physiology, National Zoological Gardens of South Africa, P.O. Box 754, Pretoria 0001, South Africa..

**Keywords:** Microclimate, evaporative cooling, biophysical ecology, operative temperature, Caprimulgus rufigena, Rufous-cheeked Nightjar

## Abstract

Nocturnally active birds roosting in exposed diurnal microsites with intense solar radiation can experience operative temperatures (T_e_) that markedly differ from air temperature (T_a_). Quantifying T_e_ thus becomes important for accurately modeling energy and water balance. We measured T_e_ at roost and nest sites used by Rufous-cheeked Nightjars (*Caprimulgus rufigena*) with three-dimensionally printed biophysical models covered with the integument and plumage of a bird. Additionally, we estimated site-specific diurnal water requirements for evaporative cooling by integrating T_e_ and T_a_ profiles with evaporative water loss (EWL) data for Rufous-cheeked Nightjars. Between 12:00 and 15:00 hrs, average T_e_ at roost sites varied from 33.1 to 49.9 °C, whereas at the single nest site T_e_ averaged 51.4 °C. Average diurnal EWL, estimated using T_e_, was as high as 10.5 and 11.3 g at roost and nest sites, respectively, estimates 3.8- and 4.0-fold greater, respectively, than when calculated with T_a_ profiles. These data illustrate that under current climatic conditions, Rufous-cheeked Nightjars can experience EWL potentially approaching their limits of dehydration tolerance. In the absence of microsite changes, climate change during the 21^st^ century could perhaps create thermal conditions under which Rufous-cheeked Nightjars exceed dehydration tolerance limits before the onset of their nocturnal active phase.

## Introduction

Organisms frequently experience the thermal environment at fine spatial scales, typically relative to their body size, resulting in microclimates that substantially differ from coarse regional macroclimates derived from standardized weather data (Beckman et al. 1973, Campbell and Norman 1998, Potter et al. 2013). An organisms immediate thermal environment arises from a complex suite of interacting abiotic variables, including solar and thermal radiation, air temperature (T_a_), wind speed, surface temperature and humidity (Porter and Gates 1969, Bakken 1976, Bakken 1989). Due to the fine scale at which microclimates occur, animals occupying the same habitat can simultaneously experience a thermally diverse range of microclimates (Sears et al. 2011), leading to large variation among individuals in energy and water demands. Behaviorally, animals may control their rate of heat loss or gain through postural changes (Porter et al. 1994) and/or by occupying thermally-buffered refugia (Wolf et al. 1996, Scheffers et al. 2014). Additionally, endotherms may temporarily abandon normothermic T_b_ by expressing patterns of thermoregulation that lead to energy and water conservation (e.g., facultative hypothermia or hyperthermia; McKechnie and Lovegrove 2002, Tieleman and Williams 1999), conditions under which microhabitat selection will have a large influence on energy and/or water savings. A thorough understanding of the microclimates an individual experiences within its habitat is thus a prerequisite for predicting energy and water requirements under current and future climates (Kearney and Porter 2009, Porter et al. 2010).

Operative temperature (T_e_), the temperature of an animal model in thermodynamic equilibrium with its environment in the absence of metabolic heating or evaporative cooling (Bakken 1976), is commonly used to quantify microclimates at spatial scales relevant to an animal (Bakken 1992, Dzialowski 2005). Operative temperature can be measured using either mathematical, statistical or biophysical models (Bakken 1992, Angilletta 2009). Biophysical models traditionally consist of a thin copper cast electroformed to match the size and shape of the focal animal and, in the case of ectotherms that lack pelages, painted to match the absorptivity of the focal species (Bakken and Gates 1975, Dzialowski 2005). For thermal investigations of endotherms, the skin and pelage of the animal of interest is typically wrapped around the cast to incorporate the thermal properties of fur or feathers (e.g., Ward and Pinshow 1995, Bozinovic et al. 2000, Tieleman and Williams 2002). Hence, biophysical models integrate the abiotic factors of the thermal environmental (i.e., radiation, T_a_, wind speed and surface temperature) with an animal’s physical attributes (i.e., size, shape and color) to determine the T_e_ experienced in a particular microsite (Bakken and Angilletta 2014).

Nightjars and nighthawks (Caprimulgidae) are a nocturnally active avian taxon that generally roost and nest on the ground during their diurnal rest phase. Several species have been reported occupying sites devoid of shade and continuously subjected to intense solar radiation, even in midsummer (e.g., Cowles and Dawson 1951, Bartholomew et al. 1962, Steyn 1971, Grant 1982, Cleere and Nurney 1998). Moreover, forced convective heat loss at these sites is likely minimal due to reduced wind speeds at ground level (Chen et al. 1998). Caprimulgids, therefore, can experience microclimates wherein T_e_ peaks at 50 - 60 °C (Weller 1958, Grant 1982, Ingels et al. 1984). Under such extreme heat, caprimulgids must elevate evaporative water loss (EWL) above baseline levels for prolonged periods to avoid lethal hyperthermia (Grant 1982, O’Connor et al. 2017b).

Our objectives were to characterize the microclimates of roost and nest sites for a southern African arid-zone caprimulgid, the Rufous-cheeked Nightjar (*Caprimulgus rufigena*). Like many other nightjars, Rufous-cheeked Nightjars may select roost and/or nest sites with partial or no shading, even in mid-summer (R.S. O’Connor personal observation, Cleere and Nurney 1998). No attempts, however, have been made to quantify the range of T_e_ values that Rufous-cheeked Nightjars experience in the field, despite the importance of these data for understanding their water budgets. We used two types of three dimensionally (3-D) printed biophysical models to measure T_e_ at roost and nest sites, one consisting of a 3-D printed plastic body (hereafter T_e-plastic_) and the second type a plastic body covered with the skin and feathers of a Rufous-cheeked Nightjar (hereafter T_e-skin_). We also integrated T_e_ and T_a_ profiles measured at each site with EWL data derived from a laboratory heat tolerance investigation for this species (O’Connor et al. 2017b) to predict site-specific water requirements for evaporative cooling during the diurnal inactive period.

## Methods

### Roost and nest location

We quantified T_e_ at six roost sites and one nest site used by six Rufous-cheeked Nightjars (two roost sites were used by the same individual) between 26 October and 12 December 2015 at Dronfield Nature Reserve (28° 39’ S, 24° 48’ E, ~1218 m a.s.l.) near Kimberley, South Africa. Nightjars were captured at night on roads with a handheld net and spotlight. Upon capture, we fitted nightjars with a radio transmitter (BD-2T, Holohil Systems, Carp, Ontario, Canada) positioned between the scapulars using a backpack-style harness built from Teflon ribbon (Telonics, Mesa, AZ, USA). Additionally, we injected a passive-integrated transponder (PIT) tag (Biomark, Boise, ID, USA) into the peritoneal cavity to register T_b_ of incubating adults as part of a separate thermoregulation study (O’Connor et al. 2017a). Birds were released immediately after transmitters were attached and PIT tags injected. We tracked birds to their roost sites using telemetry. Once a site was located, we recorded the latitude and longitude and marked the location with a metal stake for eventual placement of our models. On average, models were set up 3.5 days after discovering a roost location. For the nest site, we waited until incubation was complete before placing our models.

### Biophysical model construction and T_e_ measurements

We followed Watson and Francis (2015) and used 3-D printed models to measure T_e_. These authors compared T_e_ values of 3-D printed models to those recorded using traditional electroformed copper models and found that T_e_ distributions of the two model types were generally similar when placed in identical habitats. Furthermore, Watson and Francis (2015) reported no substantial differences in the response of models to radiant heat or varying T_a_. Finally, 3-D printed models are more anatomically accurate and easier to produce than traditional electroformed copper models, and hence are a good substitute for measuring T_e_ (Watson and Francis 2015).

We constructed four 3-D printed biophysical models, consisting of two T_e-skin_ models and two T_e-plastic_ models. To construct our T_e-plastic_ models, we took a series of 49 pictures at 90° and 45° angles encompassing 360° of a deceased Rufous-cheeked Nightjar (Supplementary Material A, Figure S1). These photos were uploaded to 123d Catch (http://www.123dapp.com/catch) which automatically stitched them into a 3-D model. Because we were unable to capture photos of the bird’s ventral surface during this procedure, we used Blender v. 2.75 (https://www.blender.org/) to manually impose a plane to the underside of each model. The digital 3-D model was then scaled using Cura (https://ultimaker.com/en/products/cura-software) and saved as an .stl file and emailed to 3DForms (Johannesburg, South Africa, http://www.3dforms.co.za/) for printing. The T_e-plastic_ models were printed using Makerbot’s (Makerbot Industries LLC, Brooklyn, NY, USA) cool grey acrylonitrile butadiene styrene (ABS) filament with a 0% fill and an approximately 2-mm thick shell. The final T_e-plastic_ models were smoothed post printing, and measured approximately 225 mm from tip of bill to tail, 44 mm from top of head to base and 53 mm wide. The T_e-plastic_ models were printed with a removable lid on the base, which we secured with adhesive prior to model deployment (Supplementary Material A, Figure S2). A type-T thermocouple (Omega, Norwalk, CT, USA) was inserted through a hole drilled in the base of each model and sealed in place with an adhesive. The tip of the thermocouple was centered horizontally and vertically within the model to avoid any effects of thermal stratification (Bakken 1992; Supplementary Material A, Figure S2). Prior to deployment, thermocouples were calibrated in a water bath between 5 and 50 °C in 5 °C increments against a mercury thermometer traceable to the US National Bureau of Standards.

To construct our T_e-skin_ models, we provided two Rufous-cheeked Nightjar carcasses to a taxidermist who separated the skin from the bodies. We took 32 pictures in series of a skinned body, again encompassing 360° and taken at 90° and 45° angles (Supplementary Material A, Figure S3). These photos were compiled, scaled and printed using the same software and plastic as outlined above. We scaled these models based on measurements provided by the taxidermist, with the final dimensions of the body measuring approximately 39.5 mm long x 14 mm wide x 23.4 mm high. Due to the smaller size of these models, they had to be printed in halves and then glued together. Consequently, some tiny gaps remained on the models and we sealed these using plaster-of-Paris and a cyanoacrylate adhesive (Supplementary Material A, Figure S4). We ensured models were airtight by completely submerging them in water to observe if any escaping air bubbles formed. We inserted a type-T thermocouple (Omega, Norwalk, CT, USA) into the approximate center of each model through a hole drilled in the ventral surface. Thermocouples were sealed in place and calibrated prior to placement as described for the T_e-plastic_ models above. The final plastic bodies were then returned to the taxidermist who wrapped them in the skin to create the complete T_e-skin_ models. Because the smoothing process left the 3-D printed plastic bodies slippery, the taxidermist had to wrap a ~5-mm layer of cotton wool around the bodies to increase adhesion between the skin and the plastic (Supplementary Material A, Figure S5).

Both T_e-skin_ and T_e-plastic_ models were placed side by side at a specific site (Supplementary Material A, Figure S6) either in the morning (mean placement time = 07:48) or at night (mean placement time = 19:12). Models were first placed facing true north, whereafter we alternated the cardinal direction approximately every 24 hours (mean time in each direction = 22.9 hours), with the models positioned in every cardinal direction before relocation to a new site. Therefore, the mean time models remained at a site was approximately 4 days (mean time at a site = 3.8 days). We recorded T_e-skin_ and T_e-plastic_ values simultaneously every minute using a 4-channel thermocouple data logger (model SD-947, Reed Instruments, Wilmington, NC, USA). We buried the thermocouple wiring between the models and logger in the sand to prevent heat from solar radiation conducting along the wire (Bakken 1992). Operative temperature data were transferred onto a personal computer every time we changed the direction of the models. Weather data were recorded every minute with a portable weather station (Vantage Pro2, Davis Instruments, Hayward, CA, USA), placed ~2.0 m above the ground and calibrated as described by Smit et al. (2013).

### Data analysis

All analyses were conducted in R v. 3.4.0 (R Core Team 2017) with values presented as mean ± standard deviation (SD). We categorized the data into three periods, namely diurnal (i.e., sunrise to sunset), midday (i.e., 12:00 - 15:00 hours) and nocturnal (i.e., sunset to sunrise). Sunrise and sunset times were calculated using the R package *maptools* (Bivand and Lewin-Koh 2017). We compared overall differences among T_e-skin_ and T_e-plastic_ models for all sites combined during the diurnal period by fitting a linear mixed-effect model using the R package *lme4* (Bates et al. 2015), with T_e_ a continuous response variable and *skin* a two-level categorical predictor. We included Te model as a random factor because of repeated T_e_ measurements within the same model. We report the effect size *skin* had on T_e_, represented as the parameter estimate (*β* ± SD) and the associated 95% confidence interval (95% CI). We considered the mean difference between T_e-skin_ and T_e-plastic_ models to be statistically significant if the 95% CI did not overlap zero. We then analyzed diel patterns in T_e-skin_ and T_a_ at each site by aggregating all values recorded each minute during a recording period and taking the average. For example, mean T_e-skin_ at 12:00 hours within a site represents the average of all T_e-skin_ values recorded at 12:00 hours at that site over the entire recording period. Because there were occasional gaps in our temperature recordings (e.g., when changing position of the models), not every minute was represented for every day and consequently sample sizes for each minute ranged from 1 to 5. To determine when and for how long T_e-skin_ exceeded free-ranging modal body temperature (T_b-mod_), we isolated all diurnal Te values > 39.7 °C at roost sites and all T_e_ values > 38.8 °C at the nest site (O’Connor et al. 2017a). We additionally calculated the direction and magnitude that T_e-skin_ deviated from free-ranging T_b-mod_ at each site during the diurnal and midday periods by subtracting T_b-mod_ from each T_e-skin_ value (i.e., ΔT_e_ - T_b-mod_). Overall roost site averages represent the combined average from each individual roost site average.

To estimate site-specific diurnal EWL during a recording period, we integrated T_e_ and T_a_ traces with the EWL data reported by O’Connor et al. (2017b). We used free-ranging T_b-mod_ for roosting birds (i.e., 39.7 °C; O’Connor et al. 2017a) as an inflection point at roost sites and the T_b-mod_ of an incubating Rufous-cheeked Nightjar (i.e., 38.8 °C; O’Connor et al. 2017a) as an inflection point at the nest site. At T_a_ and T_e_ ≤ T_b-mod_, we predicted EWL assuming EWL (g hr^−1^) = 0.007x + 0.002, and at T_a_ and T_e_ > T_b-mod_ we assumed EWL (g hr^−1^) = 0.099*x* - 3.610, where *x* represents either T_e_ or T_a_. For all T_e_ calculations we used T_e-skin_ values. We calculated a mean EWL estimate for each minute over a 24-hour period by aggregating all values at each minute as described above for the diel T_e-skin_ and T_a_ calculations. We then summed all mean EWL predictions for each minute between sunrise and sunset to obtain the total average amount of water lost during just the diurnal period at a given site. We expressed total EWL during the diurnal period as a percentage of body mass (M_b_) assuming an average M_b_ of 57.1 g, which was the average M_b_ of birds at capture when being weighed for a total body water study (O’Connor et al. *unpublished data*).

## Results

We recorded 36,648 T_e_ values for each model type. Except for a single roost situated under a camelthorn (*Vachellia erioloba*) tree, and thus shaded for most of the day, all roost sites were partially shaded and experienced periods of full solar exposure throughout the day. In contrast, the nest site was completely exposed and hence continuously subjected to intense solar radiation. Average diurnal T_a_ across sites during the study period was 28.0 ± 2.8 °C and average solar radiation level was 544.5 ± 37.3 W m^−2^. Within each site, mean T_e-skin_ values were generally similar to those of T_e-plastic_ during both the diurnal and midday periods (Table 1). On average, mean T_e-plastic_ values were 0.9 ± 2.0 °C greater than T_e-skin_ values during the diurnal period and 0.2 ± 0.6 °C greater during the midday period (Table 1). In contrast, differences between mean maximum Te within sites were larger (Table 1), with mean maximum T_e-plastic_ values 3.4 ± 3.7 °C greater than mean T_e-skin_ values. The overall difference between T_e-skin_ and T_e-plastic_ models during the diurnal period for all sites combined was not significant (skin β= -0.920 ± 3.48 °C, 95% CI = -8.14 °C, 5.93 °C), with mean T_e-skin_ = 36.3 ± 10.8 °C and mean T_e-plastic_ = 37.3 ± 11.5 °C.

**Table 1.**
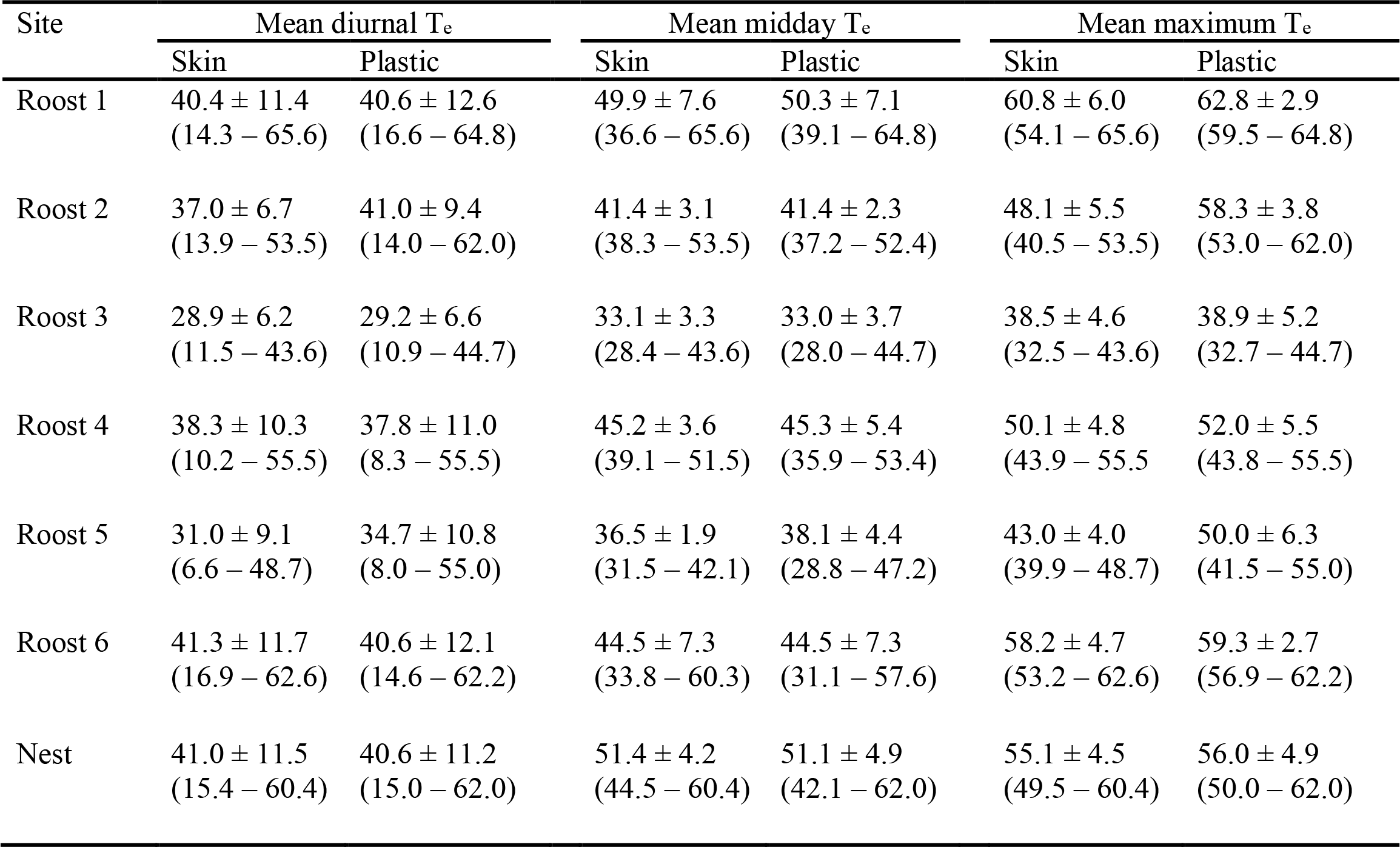
Mean ± SD diurnal (i.e., sunrise to sunset), midday (i.e., 12:00 - 15:00 hours) and maximum operative temperatures (T_e_ °C) recorded at six roost sites and one nest site used by six different Rufous-cheeked Nightjars (*Caprimulgus rufigena*; roosts 1 and 2 were occupied by the same individual). T_e_ was recorded at each site separately between 26 October and 12 December 2015, near Kimberly, South Africa at 1-minute intervals using two types of 3-D printed models, including one type covered with skin and feathers (i.e., skin) and one without skin and feathers (i.e., plastic). Sample sizes were identical between skin and plastic models, ranging from 2352 to 3497 for diurnal measurements and 543 to 806 for midday measurements. Values in parentheses represent T_e_ ranges.

During the nocturnal period, T_e_ did not deviate far from T_a_ at roost sites or the nest site (Figure 2 and Figure 3). For example, the mean difference between T_e-skin_ and T_a_ during the nocturnal period at roost and nest sites combined was 0.6 ± 2.3 °C (range = -6.8 - 8.8 °C). Beginning at sunrise, however, T_e_ increased rapidly to values far above T_a_ (Figure 2 and Figure 3). Except for at roosts 3 and 5, mean T_e-skin_ exceeded free-ranging T_b-mod_ for extended periods (Figure 2). On average, T_e-skin_ from roost sites exceeded T_b-mod_ by 08:27 h (range = 08:12 - 08:41 h; Figure 2). The overall average duration that T_e-skin_ exceeded T_b-mod_ during the diurnal period at roost sites was 6.8 hours (range = 4.6 - 7.9 hours; Figure 2). During this time, overall mean T_e-skin_ = 43.6 ± 3.5 °C (range = 39.9 - 48.9 °C) while the overall average T_a_ = 33.2 ± 3.5 °C (range = 28.1 - 38.0 °C). During the midday period, mean ΔT_e_ - T_b-mod_ among roost sites ranged from - 6.6 ± 3.3 to 10.2 ± 7.6 °C (Figure 4).

**Figure 2.**
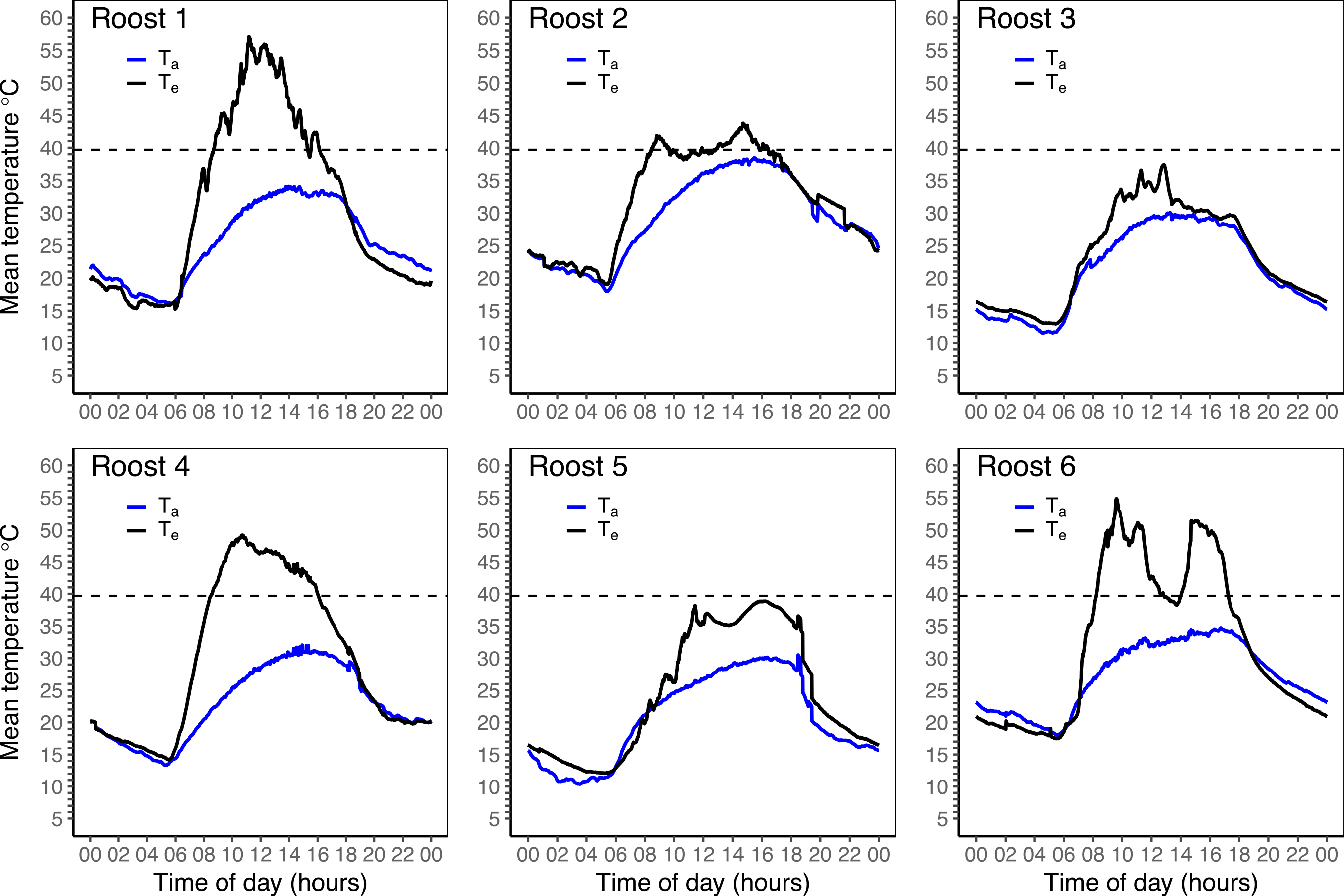
Mean operative temperature (T_e_) and air temperature (T_a_) for a 24-hour day (e.g., 02 = 02:00 hours; 22 = 22:00 hours) at six roost sites used by five Rufous-cheeked Nightjars (*Caprimulgus rufigena;* roosts 1 and 2 are from the same bird). T_e_ was recorded at 1-min intervals using a 3-D printed model covered with the skin and feathers of a Rufous-cheeked Nightjar. T_e_ was recorded at each roost site separately and remained at a site for approximately four days. Horizontal dashed lines represent free-ranging modal body temperature for roosting Rufous-cheeked Nightjars (39.7 °C; O’Connor et al. 2017a). T_a_ was also recorded at 1-minute intervals.

**Figure 3.**
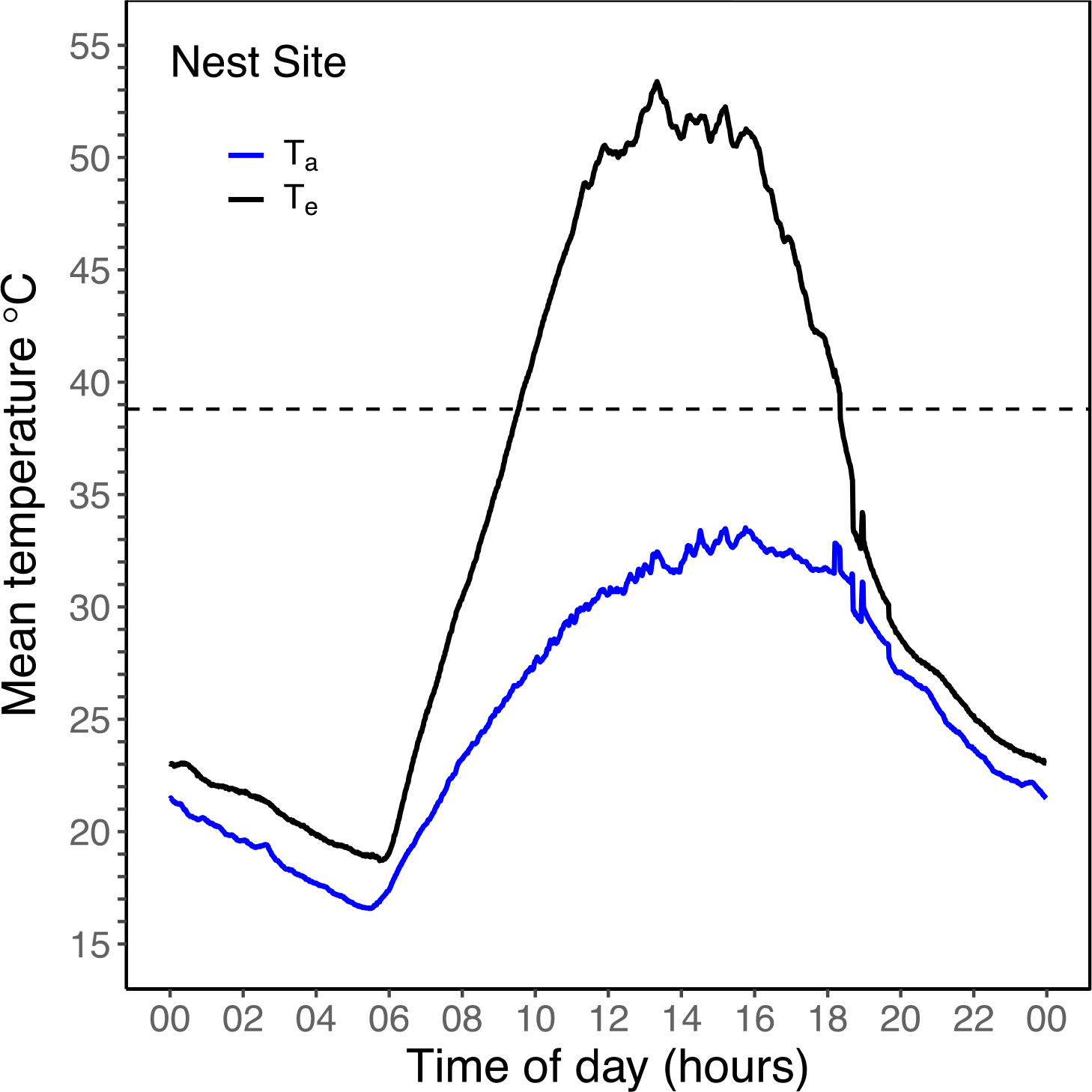
Mean operative temperature (T_e_) and air temperature (T_a_) for a 24-hour day (e.g., 02 = 02:00 hours; 22 = 22:00 hours) at a Rufous-cheeked Nightjar (*Caprimulgus rufigena*) nest site. T_e_ was recorded at 1-min intervals using a 3-D printed model covered with the skin and feathers of a Rufous-cheeked Nightjar. The model remained at the site for approximately four days. The horizontal dashed line represents the free-ranging modal body temperature for an incubating Rufous-cheeked Nightjar (38.8 °C; O’Connor et al. 2017a). T_a_ was also recorded at 1-minute intervals.

**Figure 4.**
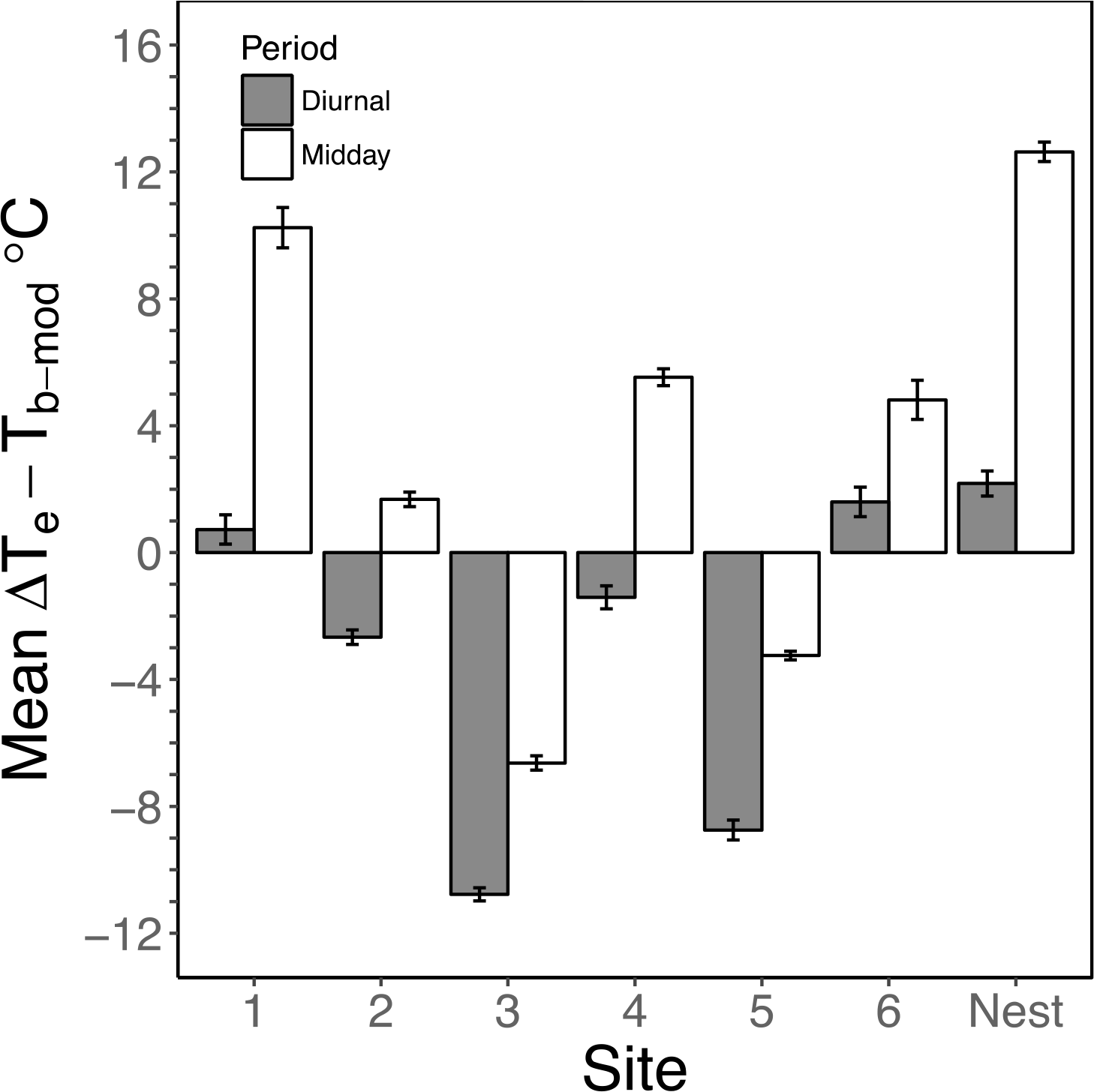
Mean difference between operative temperature (T_e_) and free-ranging modal body temperature (T_b-mod_) during the diurnal (i.e., sunrise to sunset) and midday (i.e., 12:00 - 15:00 hours) periods at six roost sites and one nest site used by six Rufous-cheeked Nightjars (*Caprimulgus rufigena*; roosts 1 and 2 are from the same individual). T_e_ was recorded at each site separately between 26 October and 12 December 2015, near Kimberly, South Africa at 1-minute intervals using a 3-D printed model covered with the skin and feathers of a Rufous-cheeked Nightjar. Error bars represent 95% confidence intervals.

Mean total diurnal EWL estimated based on T_e-skin_ ranged from 2.8 - 10.5 g among roost sites, with evaporative water requirements being 1.2 - 3.8-fold greater when calculated using T_e_ compared with estimates based on T_a_ (Figure 5). Expressed as a percentage of Mb, total diurnal EWL among roost sites ranged from 4.9 - 18.4% of M_b_.

**Figure 5.**
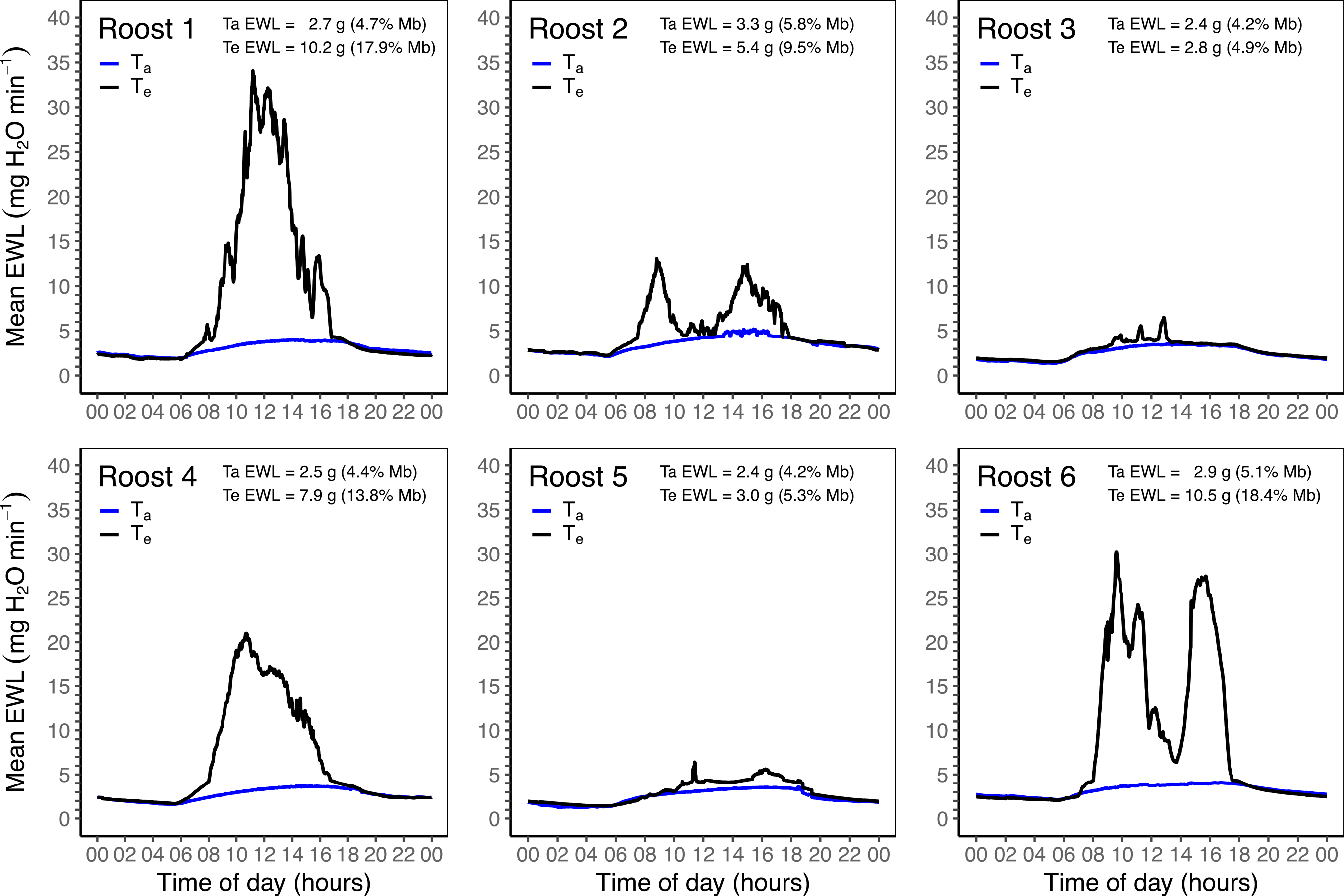
Mean evaporative water loss (EWL) predictions for a 24-hour day (e.g., 02 = 02:00 hours; 22 = 22:00 hours) at six roost sites used by five Rufous-cheeked Nightjars (*Caprimulgus rufigena*; roosts 1 and 2 were used by the same bird). EWL was predicted every minute by plugging either T_e_ or T_a_ into EWL models from O’Connor et al. (2017b). T_a_ EWL and T_e_ EWL represent the sum of all mean EWL predictions at each minute for only the diurnal period. Values in parentheses represent the amount of water lost during the diurnal period as a percentage of body mass (57.1 g).

At the nest site, mean T_e-skin_ exceeded incubating T_b-mod_ by 09:33 and did not decrease below T_b-mod_ until 18:20 (~8.75 hours; Figure 3). During this period, T_e-skin_ averaged 48.0 ± 4.0 °C (range = 38.9 - 53.4 °C), while average T_a_ = 31.3 ± 1.7 °C (range = 26.6 - 33.5 °C). The average ΔT_e_ - T_b-mod_ during the midday period at the nest site was 12.6 ± 4.2 °C (Figure 4). Diurnal evaporative water requirements at the nest site were 11.3 g when estimated from T_e-skin_, a value 4-fold greater than when estimated using T_a_ values (Figure 6). Total estimated diurnal water loss at the nest site was equivalent to 19.8% of M_b_.

**Figure 6.**
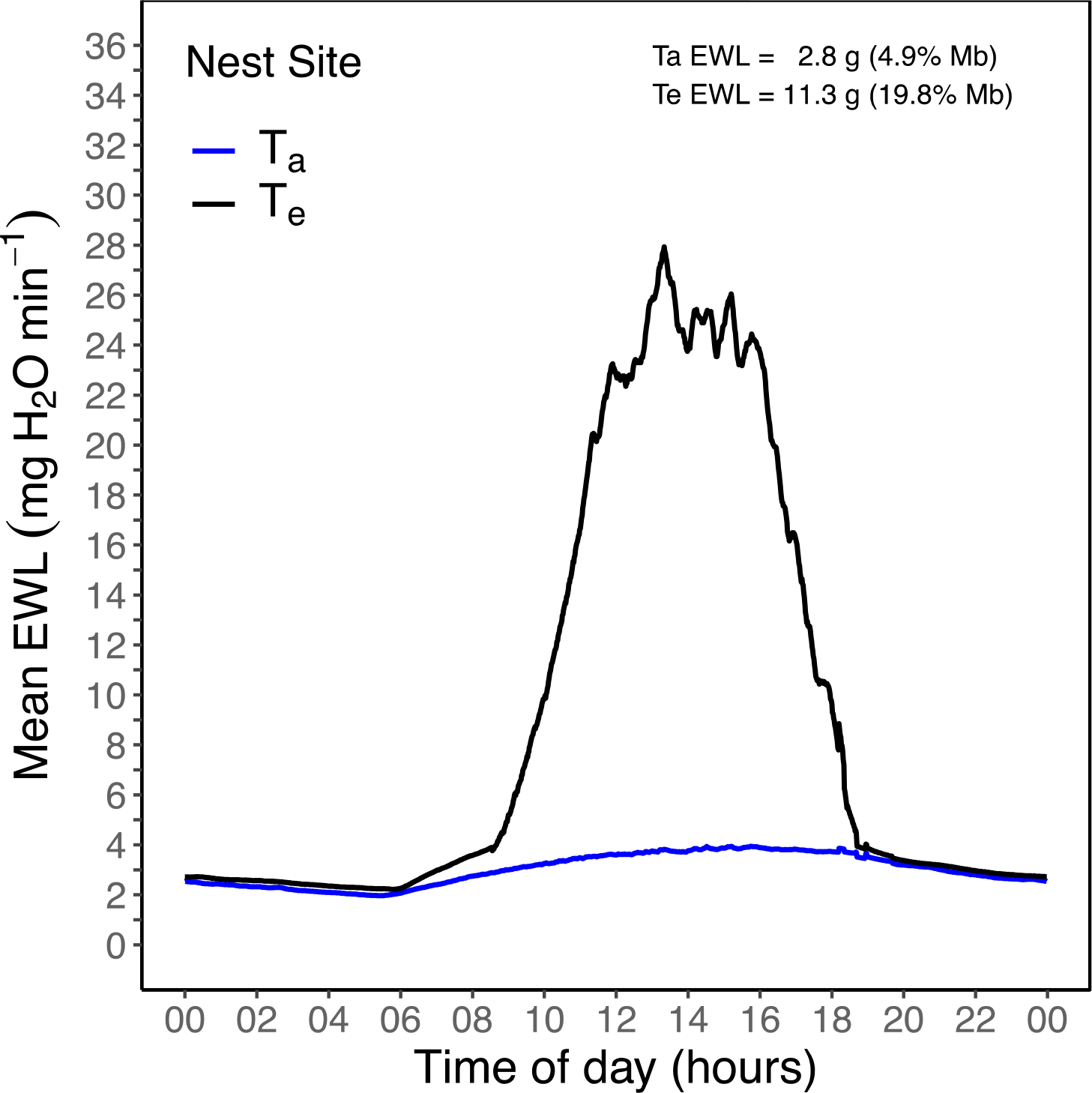
Mean evaporative water loss (EWL) predictions for a 24-hour day (e.g., 02 = 02:00 hours; 22 = 22:00 hours) for a Rufous-cheeked Nightjar (*Caprimulgus rufigena*) nest site. EWL was predicted every minute from T_e_ or T_a_ using EWL data from O’Connor et al. (2017b). Ta EWL and Te EWL represent the sum of all mean EWL predictions at each minute for only the diurnal period. Values in parentheses represent the amount of water lost during the diurnal period as a percentage of mean Rufous-cheeked Nightjar body mass (57.1 g).

## Discussion

We show that Rufous-cheeked Nightjars regularly experienced microclimates where T_e_ substantially exceeded free-ranging T_b_ (O’Connor et al 2017a). The high environmental temperatures reported here are consistent with those reported by previous authors who characterized the diurnal thermal environments occupied by nightjars and other thermally exposed birds (e.g., Weller 1958, Bartholomew and Dawson 1979, Grant 1982, Tieleman and Williams 2002, Amat and Masero 2004a, Carroll et al. 2015a). Our data further underscore the significant contribution that solar radiation can have on the total heat load of an animal and that Ta alone typically provides only a minimum index of a terrestrial animal’s thermal stress in hot environments (Porter and Gates 1969, Sears et al. 2011). This is exemplified by the fact that maximum T_a_ exceeded 38 °C on just 3 days during our recording period whereas maximum T_e_ exceeded 38 °C on 27 days.

We did not find significant differences in average diurnal T_e_ measurements among our T_e-skin_ and T_e-plastic_ models. In a similar study, Walsberg and Weathers (1986) compared T_e_ values from copper taxidermic mounts covered with the integument of four bird species to T_e_ values recorded from painted metal spheres. When averaged over a 5-day period, Walsberg and Weathers (1986) found that mean differences among the models were less than 2.0 °C. However, Walsberg and Weathers (1986) noted that when T_e_ was averaged over time scales of less than several hours, differences between models reached up to 6.3 °C. Indeed, this likely explains the larger differences we observed among our models for mean maximum T_e_ because these averages were derived from single point estimates as opposed to data spanning more than several hours. Hence, our findings support the conclusion reached by Walsberg and Weathers (1986) that complex T_e_ models may not always be necessary, as long as numerous data are collected over extended periods. Likewise, Bakken (1992) suggested that in some instances, a rough approximate representation of the study animal can be adequate and the appropriate T_e_ model used will depend on the relative importance of several considerations, such as field conditions or the study’s objective. However, investigators should attempt to use models with physical properties matching those of a study animal whenever possible (Bakken 1992, Bakken and Angilletta 2014).

Our data show that predicted diurnal water requirements can be several-fold greater when calculated using T_e_ compared to T_a_, reiterating the importance of using spatially relevant microclimates when assessing an animal’s physiological stress (Huey 1991, Helmuth et al. 2010, Porter et al. 2010). Moreover, the amount of water lost based on T_e_ values was equivalent to a substantial percentage of M_b_, suggesting that, on hot, cloudless days, Rufous-cheeked Nightjars might be approaching their limits of dehydration tolerance. Unfortunately, few data exist on acute dehydration tolerance among birds when exposed to severe heat stress over time scales of hours on very hot days (Wolf 2000, McKechnie and Wolf 2010, Albright et al. 2017). Wolf and Walsberg (1996), however, reported acute dehydration tolerance limits in Verdins (*Auriparus flaviceps*; ~7 g) when water loss exceeded 11% of M_b_, a physiological threshold far lower than the maximum water losses predicted here. An important factor known to enhance dehydration tolerance among birds and mammals is the ability to conserve plasma volume (Horowitz and Borut 1970, Arad et al. 1989, Carmi et al. 1993). However, plasma volume conservation is apparently affected by the Ta at which dehydration occurs (Carmi et al. 1994). Carmi et al. (1994), for example, found that Rock Pigeons (*Columbia livia*) could maintain plasma volume at T_a_ of 36 °C for ~32 hours but, when exposed to T_a_ of 40 °C for ~28 hours, plasma volume decreased by 8.9%, despite similar total losses in water between the T_a_ groups. To our knowledge, there are no data on whether caprimulgids conserve plasma volume during acute heat stress, but we speculate that Rufous-cheeked Nightjars and relatives have evolved mechanisms increasing permeability for osmotic diffusion between the extracellular and intracellular compartments, thereby aiding plasma volume conservation and allowing them to tolerate prolonged periods of high EWL.

Despite a large rapid depletion of body water during the diurnal rest phase, we predict that Rufous-cheeked Nightjars with a mean M_b_ of 57.1 g can periodically replenish body water by obtaining a maximum of 10.3 g H_2_O through preformed and metabolic water (Supplementary Material B). However, the temporal window for nightjars to forage is highly variable and constrained by several ecological and environmental factors (Mills 1986, Jetz et al. 2003, Ashdown and McKechnie 2008, Woods and Brigham 2008). Consequently, nightjars may not always acquire enough insects to offset EWL, increasing their dependence on drinking water. Indeed, on multiple occasions at dusk we observed Rufous-cheeked Nightjars drinking on the wing at water reservoirs near roost and nest sites. Furthermore, assuming a proportional increase in T_e_ with the projected 4 °C increase in T_a_ during the 21^st^ century (Smith et al. 2011), maximum predicted water requirements at roost and nest sites for Rufous-cheeked Nightjars at Dronfield could reach values of 13.8 and 15.1 g, respectively, equivalent to 24.2 and 26.4% of M_b_. Presumably, nightjar populations with no access to drinking water will be highly vulnerable to climate change because of the increasing difficulty of offsetting water deficits solely through preformed and metabolic water.

Given the design and construction of our T_e_ models, it is possible that multiple sources of error were introduced in our T_e_ estimates. Firstly, the size difference between our 3-D printed bodies could have created issues with thermal stratification (Bakken 1992, O’Connor et al. 2000, Bakken and Angilletta 2014). However, the generally small size of our models (< 100 g) combined with the thickness of the plastic (~2 mm) and the placement of the thermocouples likely mitigated this issue (Bakken 1992). The second potential source of error stems from the layer of cotton wool wrapped around the plastic body of our T_e-skin_ models. This cotton added an insulative layer which likely increased the time constant of our models. Bakken (1992) and O’Connor (2000) proposed that the time constant desired ultimately depends on the study question and rapid time responses are plausibly more important in studies on animals where behavioral thermoregulation is paramount (e.g., ectotherms). Because nightjars are inactive during the day, behavioral thermoregulation is minimal, aside from postural adjustments, and an instantaneous time constant may not be as imperative. In any event, because we lacked the necessary equipment (e.g., wind tunnel and solar simulator), we could not accurately calibrate our models prior to use, an issue that appears to be common among operative temperature studies (Walsberg and Wolf 1996, Dzialowski 2005).

## Conclusions

One of the most pressing issues facing biologists today is predicting how organisms will respond to climate change (Schwenk et al. 2009, Sears and Angilletta 2011). A vital step towards addressing this issue is knowing the degree to which an organism is exposed to environmental change (Williams et al. 2008). Organismal exposure, however, will be mediated through microhabitat selection and the use of microrefugia which can substantially buffer or amplify an environmental signal (e.g., Woods et al. 2015, Morelli et al. 2016, Pincebourde et al. 2016, Lenoir et al. 2017). Hence, organisms usually experience microclimates at spatial scales much finer than those recorded at gridded weather stations (Campbell and Norman 1998, Helmuth et al. 2010). An understanding of the thermal heterogeneity an organism is exposed to across its range of microclimates is necessary when assessing its physiological stress under current and future climate conditions.

During our study, Rufous-cheeked Nightjars experienced microclimates where T_e_ substantially exceeded normothermic T_b_ for periods of several hours each day. Although diurnally active birds may also experience similarly high environmental temperatures when foraging, they also can periodically escape midday heat by seeking out shaded microhabitats with more moderate microclimates (Goldstein 1984, Carroll et al. 2015b, Pattinson and Smit 2017). Hence, diurnal birds can reduce rates of EWL through behavioral thermoregulation (Williams et al. 1999, Wolf 2000). In contrast, nightjars remain inactive and experience the full brunt of the sun, resulting in large evaporative water requirements (Grant 1982). The capacity for nightjars to tolerate high heat loads stems from a combination of a low resting metabolic rate and an energetically efficient mechanism for dissipating heat (Dawson and Fisher 1969, O’Connor et al. 2017b, Talbot et al. 2017). Both of these traits serve to reduce an individual’s total heat load by minimizing endogenous heat production. Several authors have suggested that the use of exposed sites by ground nesting species presents a trade-off between lower predation risk due to early predator detection and increased heat stress (Amat and Masero 2004b, Tieleman et al. 2008). However, increasing temperatures could alter the dynamics of this trade-off, possibly forcing nightjars to use more shaded microhabitats despite greater predation risk. Our EWL estimates based on T_e_ values suggest that Rufous-cheeked Nightjars require exceedingly high EWL rates. The physiological mechanisms allowing nightjars to tolerate large losses in body water are presently unknown, but could pertain to an increased capacity to maintain plasma volume. Additionally, free-ranging Rufous-cheeked Nightjars likely conserve water through facultative hyperthermia (O’Connor et al. 2017a). Under current thermal conditions, Rufous-cheeked Nightjars can apparently offset evaporative water losses with preformed water from their diet and metabolic water. Climate change, however, will likely increase evaporative water requirements, in turn increasing the importance of drinking water and, possibly, even resulting in conditions where nightjars cannot make it to sunset without becoming lethally dehydrated.

## Acknowledgements

We sincerely thank Alex Mullinos at 3dforms for his patience and help with 3-D printing and Dr. James Meyer for performing the taxidermy necessary to complete the models. Without their assistance, this study would not have been possible. We additionally thank Duncan MacFadyen and E. Oppenheimer & Son for granting us access to their property. Cathy Bester and all field assistants provided invaluable help during the field season. Lastly, Bruce Woodroffe and Awesome Tools (Cape Town, South Africa) provided discounted lighting equipment, for which we are greatly appreciative.

